# PHi-C: deciphering Hi-C data into polymer dynamics

**DOI:** 10.1101/574962

**Authors:** Soya Shinkai, Masaki Nakagawa, Takeshi Sugawara, Yuichi Togashi, Hiroshi Ochiai, Ryuichiro Nakato, Yuichi Taniguchi, Shuichi Onami

## Abstract

Computational modelling methods for Hi-C data have revealed averaged and static features of the 3D genome in cell nuclei. Here, we describe a 4D simulation method, PHi-C (Polymer dynamics deciphered from Hi-C data), that depicts dynamic 3D genome features through polymer modelling. This method allows for demonstrations of dynamic characteristics of genomic loci and chromosomes, as observed in live-cell imaging experiments, and provides physical insights into Hi-C data.

---

Genomes consist of one-dimensional DNA sequences and are spatio-temporally organized within the cell nucleus. Contact frequencies in the form of matrix data, measured using genome-wide chromosome conformation capture (Hi-C) technologies, have uncovered three-dimensional (3D) features of average genome organization in a cell population ^1, 2^. Moreover, live-cell imaging experiments can reveal dynamic chromatin organization in response to biological perturbations within single cells ^3, 4^. Bridging the gap between these different sets of data derived from population and single cells is a challenge for modelling dynamic genome organization ^5, 6^.

Several modelling methods have been developed to reconstruct 3D genome structures and predict Hi-C data ^7, 8^. In addition, there has been development of bioinformatic normalization techniques in Hi-C matrix data processing to reduce experimental biases ^9–11^. However, the meaning of a contact matrix as quantitative probability data has not been discussed; moreover, a four-dimensional (4D) simulation method to explore dynamic 3D genome organization remains lacking.

Here, we introduce PHi-C, a method that can overcome these challenges by polymer modelling from a mathematical perspective and at low computational cost. PHi-C is a method that deciphers Hi-C data into polymer dynamics simulations (Fig. 1a, https://github.com/soyashinkai/PHi-C). PHi-C uses Hi-C contact matrix data generated from a *hic* file through JUICER ^12^ as input (Supplementary Fig. 1a). PHi-C assumes that a genomic region of interest at an appropriate resolution can be modelled using a polymer network model, in which one monomer corresponds to the genomic bin size of the contact matrix data with attractive and repulsive interaction parameters between all pairs of monomers described as matrix data (Methods, Supplementary Note). Instead of finding optimized 3D conformations, we can utilize the optimization procedure (Supplementary Fig. 1b,c) to obtain optimal interaction parameters of the polymer network model by using an analytical relationship between the parameters and the contact matrix. We can then reconstruct an optimized contact matrix validated by input Hi-C matrix data using Pearson’s correlation *r*. Finally, we can perform polymer dynamics simulations of the polymer network model equipped with the optimal interaction parameters.

**Figure 1:**
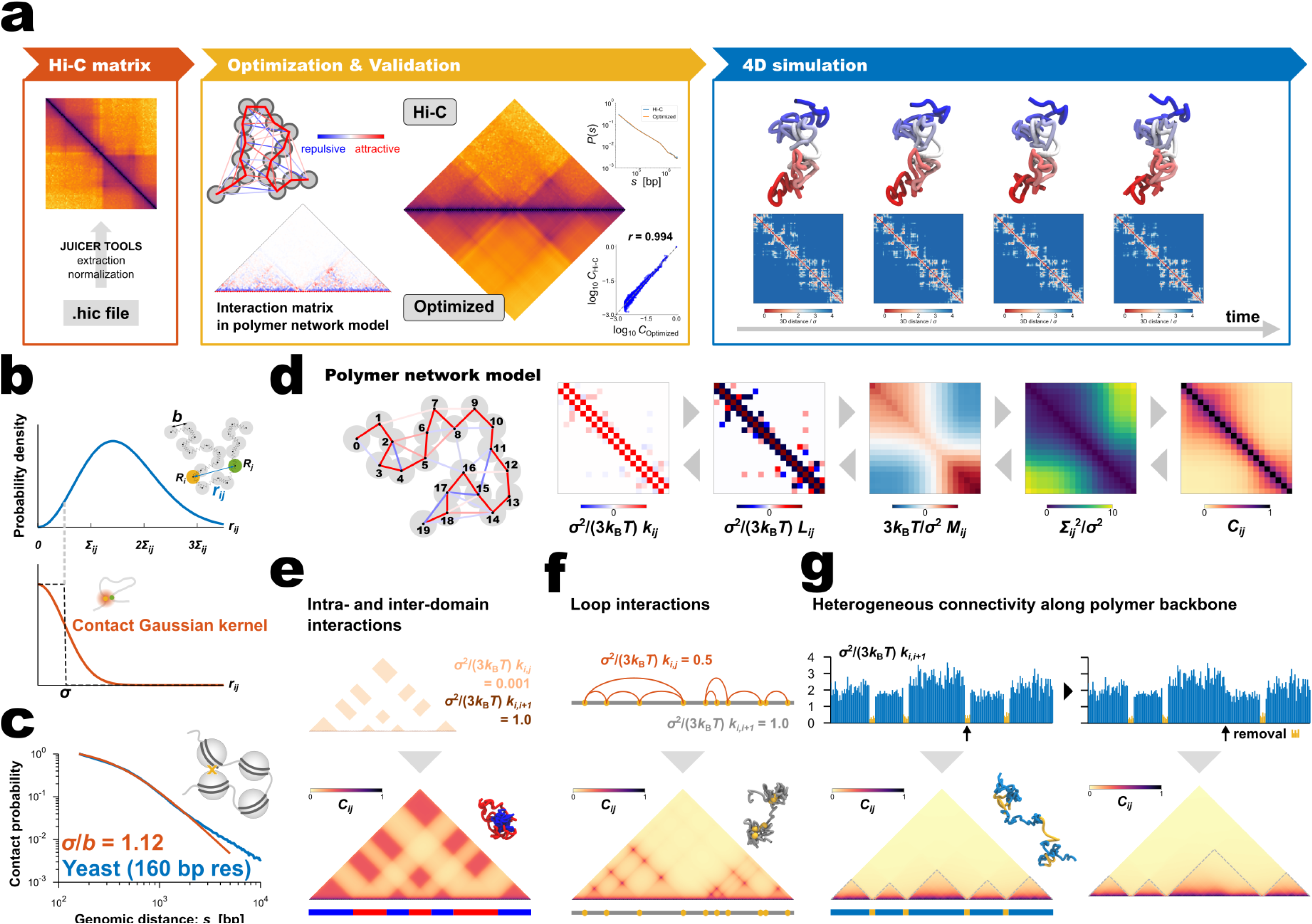
Features of PHi-C. a, Overview of PHi-C procedure. **b**, (Upper) In the bead-spring model, the probability density of the distance *r*_*ij*_ between the *i*-th and *j*-th beads (or monomers) is characterized by only the standard deviation S_*ij*_. (Lower) The contact Gaussian kernel function can capture contacts with the contact distance *σ*. **c**, Contact probability (blue) for yeast cells with 160-bp nucleosome-resolution ^14^ as a function of genomic distance averaged across the genome, and the theoretically fitted curve (orange) at a small genomic distance. **d**, The polymer network model is characterized by connectivity between all pairs of monomers, expressed by the interaction matrix *k*_*ij*_. The matrix *k*_*ij*_ is reversibly converted into the contact matrix *C*_*ij*_ through matrix transformations. Each matrix has dimensionless values with a normalization factor. **e–g**, Painting contact patterns for intra- and inter-domain interactions (**e**), loop interactions (**f**), and heterogeneous connectivity along the polymer backbone (**g**). (Upper) Designed interactions in the polymer network model. (Lower) Converted contact matrix, with a snapshot of polymer conformations in the polymer dynamics simulation (Supplementary Videos 1–3). **g**, TAD-like domains are highlighted by dashed lines. (Right) Removal of a domain-boundary part (yellow) results in domain fusion.

First, we evaluated PHi-C’s theoretical assumption about chromosome contact. Here, we started with a simple polymer model called the bead-spring model, in which the characteristic length *b* between adjacent beads (or monomers) represents the physical size corresponding to one genomic bin of the contact matrix data. To mathematically define the contact between a pair of monomers, we introduced the contact Gaussian kernel with the contact distance *σ* (Fig. 1b). The above assumption can be used to derive the theoretical scaling relationship of the contact probability, 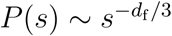, as a function of genomic distance *s* in terms of the fractal dimension of polymer organization *d*_f_ ^13^ (Supplementary Fig. 2a,b). In addition, interestingly, the ratio of the contact distance to the length between adjacent monomers, *σ*/*b*, makes the shape of the contact probability rounder at a small genomic distance (Supplementary Fig. 2b). This phenomenon implies that the rounded shape conveys information about the ratio *σ*/*b*. To assess our theoretical framework about contact and how the contact distance varies in Hi-C experiments, we analysed yeast Hi-C data at nucleosome resolution ^14^. The fitted value was *σ*/*b* = 1.12, suggesting that contacts mainly occur within a distance corresponding to the size of a nucleosome in this super-resolution Hi-C experiment (Fig. 1c, Supplementary Fig. 2c). Other high-resolution Hi-C data for human GM12878 ^10^ revealed *σ*/*b* = 1.38, suggesting that cross-linking of Hi-C experiments almost exactly captures chromosome contacts with an appropriate resolution (Supplementary Fig. 2d).

**Figure 2:**
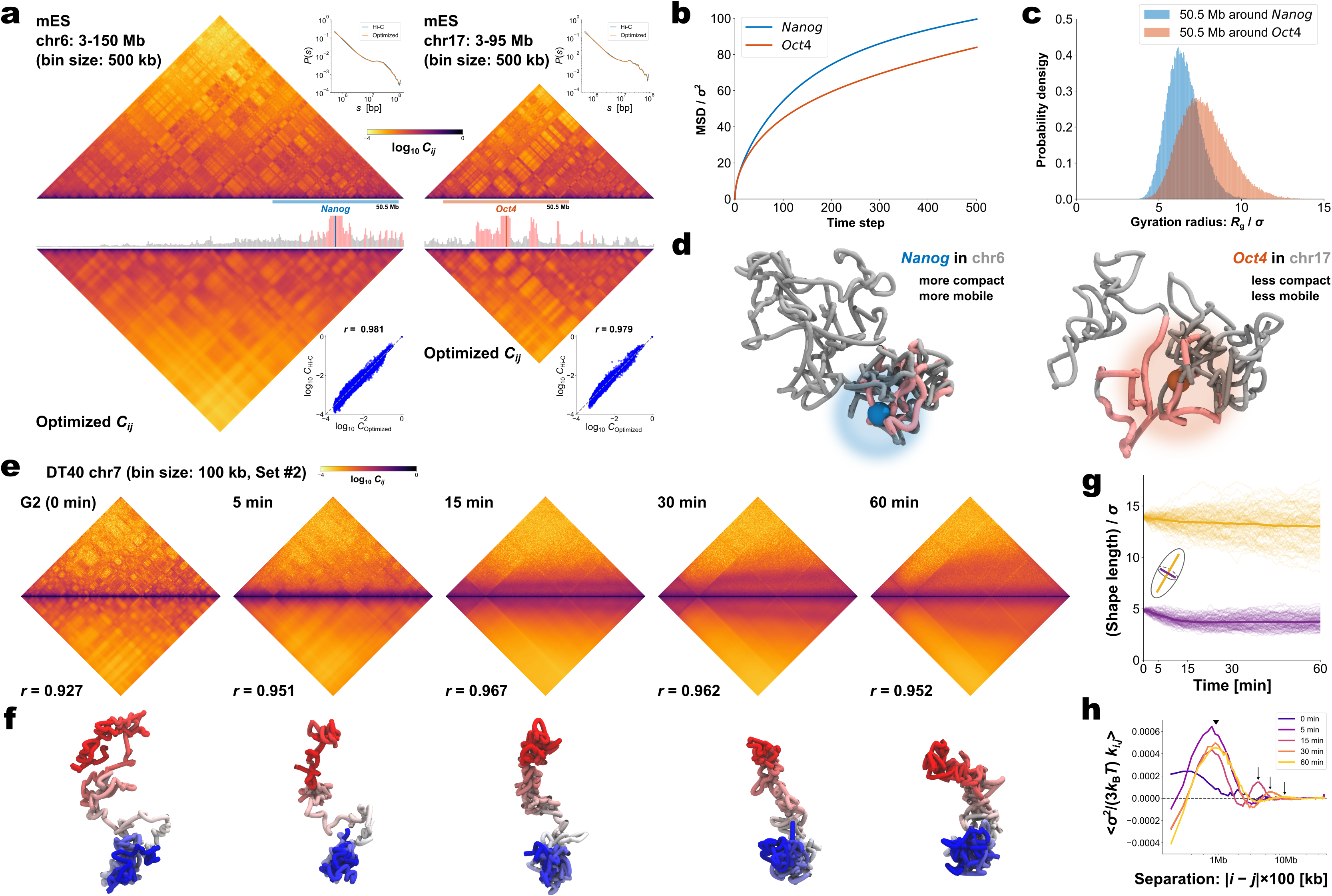
Demonstrations of PHi-C for Hi-C data of mESCs and DT-40 cells. a, PHi-C analysis for chromosomes 6 (left) and 17 (right) of mESCs. (Upper) Contact matrices of the Hi-C experiment (binned at 500 kb), and contact probabilities as a function of genomic distance. (Middle) 4C-like profiles of *Nanog* (left) and *Oct4* (right) loci. High-interaction regions are highlighted (pink). (Lower) Optimized contact matrices by PHi-C, and correlation plots between log_10_ C_Hi*-*C_ and log_10_ C_Optimized_. **b**, Theoretical MSD curves of *Nanog* and *Oct4* loci. **c**, Probability densities of the gyration radius of 10^5^ conformations for the 50.5-Mb genomic regions around *Nanog* and *Oct4* loci in mESCs. **d**, Polymer models derived from PHi-C analysis for *Nanog* and *Oct4* loci on chromosomes 6 and 17, respectively. Pink-highlighted regions on the polymer models correspond to the regions in the 4C-like profile of **a**. **e**, PHi-C analysis for chromosome 7 of DT-40 cells at G2 (0 min), 5, 15, 30 and 60 min. Contact matrices of the Hi-C experiment with 100-kb-sized bins (upper) and optimized contact matrices with the correlation value (lower). **f**, Snapshots of polymer conformations in a 4D polymer dynamics simulation. **g**, Time series of the shape lengths of the major (yellow) and minor (purple) axes for polymer conformations in 100 polymer dynamics simulations starting from the same initial conformation. Thick curves represent the averages. **h**, Curves of optimized interaction parameters, 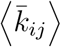, averaged at each genomic distance (separation, |*i-j*| *×* 100 kb). A triangle indicates a position of a local peak inducing compaction within 2 Mb. Arrows indicate positions of a local peak generating periodicity of attractive interactions around 4, 6 and 10 Mb at 15, 30 and 60 min, respectively.

An important step in the optimization procedure is based on analytical matrix transformations between the polymer network model and the contact matrix (Fig. 1d, Methods). The matrix transformations provided us with a low computational cost optimization strategy that can be applied to find optimal interaction parameters of the polymer network model without sampling optimal static 3D polymer conformations (Supplementary Fig. 1b,c). Moreover, we can depict any contact patterns in a moment by using the matrix transformations and perform polymer dynamics simulations by designing interactions in the polymer network model (Fig. 1e–g, Supplementary Videos 1–3). Intra- and inter-domain interactions generate a chequerboard pattern reminiscent of A/B compartments ^1, 2^, and the attractive interaction domains form a combined domain (Fig. 1e). Loop interactions show a clear punctate pattern (Fig. 1f). Furthermore, we can depict a topologically associating domain (TAD)-like pattern ^1, 2^ by only tuning heterogeneous connectivity along the polymer backbone, where less connected regions behave as domain boundaries (Fig. 1g; left). The 3D conformation suggests that the boundary regions are physically elongated with insulating inter-TAD-like-domain interactions. In addition, the removal of a boundary part causes the adjacent domain fusion (Fig. 1g; right) that reminds us of fusions of TADs ^8, 15^.

To investigate how PHi-C explains 4D features of chromosomes within living cells, we applied this approach to Hi-C data for mouse embryonic stem cells (mESCs)^16^. A live-cell imaging experiment showed a marked difference in the movements of *Nanog* and *Oct4* loci in mESCs ^17^: statistically significant enhancement of *Nanog* diffusive movement compared with *Oct4* diffusive movement was revealed (Supplementary Fig. 3). The optimization step of the PHi-C analysis for chromosomes 6 and 17 provided optimized contact matrices with correlations of more than 97% between the Hi-C and optimized contact matrices (Fig. 2a). The mean-squared displacement (MSD) curves that were theoretically derived from the optimized data for the *Nanog* locus on chromosome 6 and the *Oct4* locus on chromosome 17 are consistent with the experimental dynamics (Fig. 2b). We also compared the physical sizes of 50.5-Mb genomic regions around the *Nanog* and *Oct4* loci, with the inclusion of several areas that highly interact with each locus, and observed more compact organization of the *Nanog* region (Fig. 2c). Taking together, these findings indicate that PHi-C analysis can provide new insights into genome organization and dynamics: for example, a region of 50.5 Mb around *Nanog* adopts a more compact organization than an equivalent region around *Oct4*, and the *Nanog* locus on chromosome 6 is more mobile than the *Oct4* locus on chromosome 17 (Fig. 2d).

Finally, we used PHi-C to demonstrate the dynamic chromosome condensation process for the highly synchronous entry of DT-40 cells, which revealed a pathway for mitotic chromosome formation ^18^. The optimized contact matrices at five different time points were reconstructed with high correlations (Fig. 2e, Supplementary Fig. 4). Using the optimized interaction parameters in the polymer network model (Supplementary Fig. 5), we conducted 4D simulations starting from a comparatively elongated conformation at 0 min. The polymer conformation dynamically changed into a rod-shaped structure, revealing the condensation state of chromosomes at prometaphase (Fig. 2f, Supplementary Videos 4, 5). We evaluated dynamic changes in polymer conformations in simulations by calculating the characteristic shape lengths of the major and minor axes. As observed by microscopy, rapid and gradual decreases in the minor and major axes within 15 and 60 min indicated thin and thick rod-shaped formations, respectively (Fig. 2g). In addition, the optimized interaction parameters averaged at each genomic separation represent not only strong compaction within 2 Mb during mitosis but also increase of periodicity of long-range attractive interactions from 3 Mb to 12 Mb in prometaphase (Fig. 2h). These physical findings are consistent with a helical organization in rod-shaped chromosomes during prometaphase ^18^.

We have shown that PHi-C can decipher Hi-C data into polymer dynamics, based on a mathematical theory of chromosome contacts in the polymer network model. As shown for mESC Hi-C data, PHi-C analysis can bridge the gap between Hi-C data and imaging data with respect to chromatin dynamics in living cells. In addition, PHi-C’s theoretical basis allows for the depiction of any Hi-C pattern by designing the appropriate interaction parameters, which supports a model for TAD formation: physical chromatin stiffness based on certain molecular interactions creates insulation at TAD boundaries ^19^. Polymer modelling studies have revealed that chromatin modifications alter the physical properties of chromatin fibres and affect chromosome organization ^11, 20^. Because PHi-C analysis can extract physical interaction parameters from Hi-C data, it should be elucidated which molecular interactions on chromatin are related to physical parameters. Further comprehensive PHi-C analysis could provide physical insights into molecular interactions on chromosomes.

## Acknowledgments

We thank K. Shiroguchi and K. Kinoshita for helpful comments. We thank M. Marti-Renom and M. Di Stefano for stimulative discussions. We thank I. Hiratani for critical reading of this manuscript. We also thank Nature Research Editing Service for English language editing. This work was supported by JSPS KAKENHI Grant Numbers 16H01408 and 18H04720 (to S.S.).

## Author Contributions

S.S., M.N. and T.S. conceived the mathematical concept to define the chromosome contacts. S.S. and S.O. designed the study. S.O. supervised the study. S.S., M.N. and T.S. performed mathematical calculations. S.S. and Y.To. developed the optimization algorithm. S.S. and Y.To. wrote the codes. S.S., R.N. and Y.Ta. analysed the Hi-C data. S.S. and H.O. analysed the movements of genome loci. S.S. wrote the manuscript.

## Methods

### Overview of PHi-C

PHi-C enables us to decipher Hi-C data into polymer dynamics simulation. PHi-C is based on a theory of the polymer network model and defining contacts between two monomers on the polymer (Supplementary Note). The theoretical framework provides the following matrix transformations (Fig. 1d), where the matrix size is *N* × *N*: (i) the normalized interaction matrix 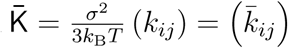 into the normalized Laplacian matrix 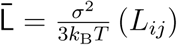 by 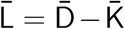, where the normalized degree matrix 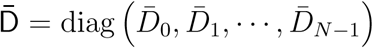, where 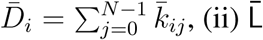 into the normalized covariant matrix relative to the centre-of-mass 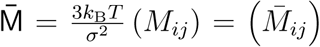 by 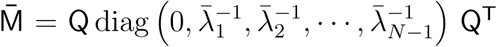, where 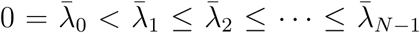 are eigenvalues of the matrix 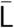 and Q is the orthogonal matrix satisfying 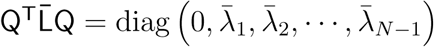,(iii) 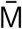 into the normalized variance matrix 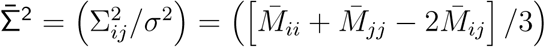, and (iv) 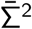 into the contact matrix 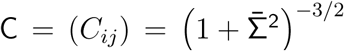. The matrix 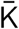 describing attractive and repulsive interactions of the polymer network model is optimized so that the difference between an input Hi-C contact matrix C_Hi*-*C_ and the reconstructed contact matrix C_reconstructed_, through the above transformations, is minimized. Finally, a 4D simulation of the polymer network model with the optimized matrix 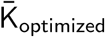 is performed. Below, we describe each step in Fig. 1a in detail.

#### Input data

PHi-C requires *N × N* contact matrix data for a genomic region of interest as an input, which is generated through the JUICER and JUICER TOOLS ^12^ from public Hi-C data with a normalization option (VC / VC SQRT / KR). Here, we used the KR normalization ^21^. In our theoretical framework, the diagonal elements of the contact matrix should satisfy *C*_*ii*_ = 1, so we additionally normalized the contact matrix such that the shape of the contact probability as a function of genomic distance *P* (*s*) is unaltered, with an interpolation if needed (Supplementary Fig. 1a). Note that the interpolation may result in artefacts for sparse contact matrix data. Finally, we obtained a normalized Hi-C contact matrix C_Hi*-*C_.

#### Optimization and validation

The optimization algorithm is designed to minimize the Frobenius norm ‖ log_10_ C_reconstructed_ *-* log_10_ C_Hi*-*C_‖_F_ as a cost function, where the contact matrix C_reconstructed_ is generated from the normalized interaction matrix 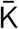. At every optimization step, an integer pair (*i, j*) is randomly selected, and the values of 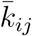 and 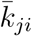 are slightly altered. If the alteration decreases the cost function, the matrix 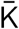 is updated. A flowchart of the algorithm is presented in Supplementary Fig. 1b.

As our optimization method is based on the random sampling of integer pairs, the amount of calculation in the procedure is proportional to *O* (*N* ^2^). In our demo codes, the hyper-parameters for the optimization are tuned, and it takes about 13 min to obtain an optimized solution for *N* = 97, even on our laptop PC (Intel^®^ CoreTM i7-6600U, dual-core 2.60GHz).

After optimization, an optimized contact matrix C_optimized_ is converted from an optimized matrix 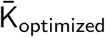. To assess the compatibility between contact matrices C_optimized_ and C_Hi*-*C_ in a logarithmic scale, we used Pearson’s correlation coefficient.

#### Polymer dynamics simulation

We performed 4D simulations of the polymer network model by using the normalized interaction matrix 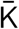. First, 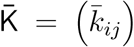 is converted into the normalized Laplacian matrix 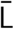. Using the eigendecomposition of the matrix 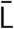, the normalized eigen-values 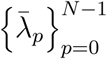 and the orthogonal matrix Q are obtained. For a normalized polymer conformation vector 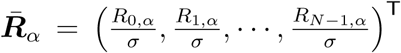, where *R*_*i,α*_ stands for the *α* (= *x, y, z*) coordinate of the *i*-the monomer, the converted vector 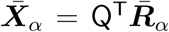 satisfies the variance relationship 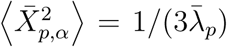 for *p* = 1, 2, *…, N -* 1. Therefore, an initial conformation of the converted vector in thermal equilibrium is given: 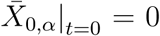, so that the centre-of-mass is the origin, and 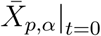 is a random variable obeying the normal distribution with mean 0 and variance 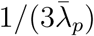 for *p* = 1, 2, *…, N -*1. Then, the initial normalized conformation in thermal equilibrium is calculated as 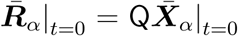. Finally, to calculate the polymer dynamics, we numerically integrated the stochastic differential equation (SDE) by using Heun’s method ^22^: the integral algorithm is defined by first predicting

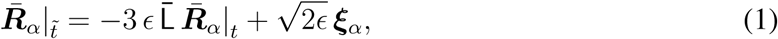

and then correcting

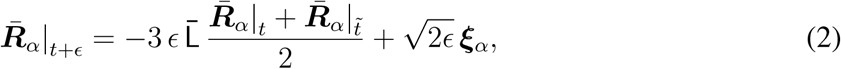

where 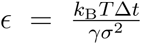 and the vector ***ξ***_*α*_ = (*ξ*_0,*α*_, *ξ*_1,*α*_, *…, ξ*_*N-*1,*α*_)^T^ consists of random variables 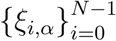 obeying the normal distribution with mean 0 and variance 1. The parameter *ϵ* is non-dimensional and represents a normalized step time, and it determines the accuracy of the SDE integration. Δ*t* stands for the step time of the integration in actual time. Here, we set *ϵ* = 0.0001.

#### Visualization of polymer conformation

The code for polymer dynamics simulation outputs XYZ and PSF files to visualize the simulated polymer dynamics. Polymer conformations were visualized using VMD ^23^ by reading these files.

### Fitting contact probability

We theoretically derived how the contact probability as a function of genomic distance averaged across the genome, *P* (*s*), behaves in terms of the fractal polymer (Supplementary Note, Supplementary Fig. 2). The function from high-resolution Hi-C data was fitted by

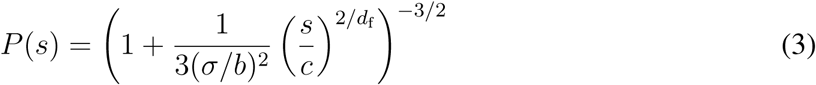

for a small genomic region and

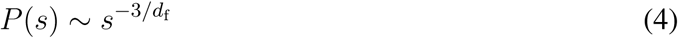

for a large genomic region. Here, the ratio *σ*/*b* and the fractal dimension *d*_f_ are the fitted parameters, and *c* stands for the genomic size corresponding to the bin size of the Hi-C matrix. We used the nonlinear least-squares Marquardt–Levenberg algorithm on GNUPLOT.

### Calculating MSD of genome loci

We re-analysed movements of *Nanog* and *Oct4* loci in mESCs. In each session of live imaging, 3D time-series of a genome locus (*Nanog* or *Oct4*) and the nucleus centre-of-mass, 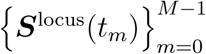 and 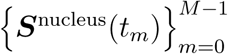, were simultaneously acquired ^17^, where the maximum frame number was *M* = 50, the time interval was Δ*t* = 10 s, and *t*_*m*_ = *m* Δ*t* (*m* = 0, 1, 2, *…, M -* 1). To eliminate the effect of movement of the nucleus, we dealt with movements of the locus relative to the nucleus centre described by the time-series 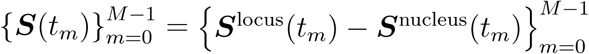. Then, the time-averaged mean square displacement (TAMSD) for a time-series 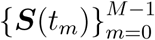 was calculated as follows:

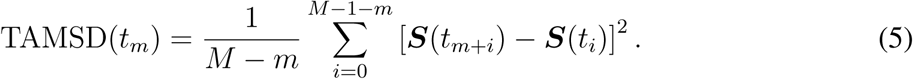

### Calculating theoretical MSD curve

The optimized matrix 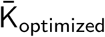 derives a theoretical MSD curve for the *i*-th monomer in the polymer network model as follows (Supplementary Note):

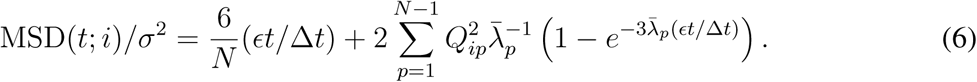

Here, not only is the MSD normalized by *σ*^2^ in the length scale, but the time-step is also normalized in time, that is, MSD*/σ*^2^ and *ϵt/*Δ*t* are dimensionless.

### Calculating radius of gyration

As we described in *polymer dynamics simulation*, a normalized polymer conformation 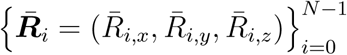 in thermal equilibrium can be sampled on the basis of the optimized matrix 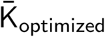. We calculated the radius of gyration for the 50.5-Mb genomic regions around *Nanog* and *Oct4* loci in mESCs. By using two integers *n*_start_ and *n*_end_ corresponding to the 50.5-Mb region, the radius of gyration is calculated as

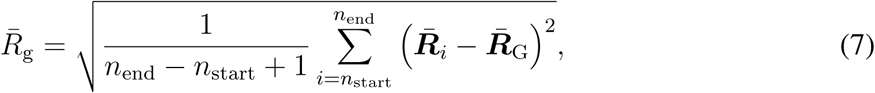

where 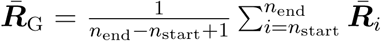 represents the centre-of-mass of the polymer conformation of the 50.5-Mb genomic region.

### Simulating polymer dynamics during chromosome condensation

We applied PHi-C to Hi-C data during mitotic chromosome formation in chicken DT-40 cells ^18^. We used the second dataset of chromosome 7 (binned at 100 kb) for the wild type at G2 (0 min), 5, 15, 30 and 60 min. We eliminated the centromere region due to the lack of associated read counts. Through the optimization of PHi-C, we obtained the optimized matrices 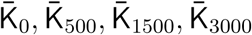 and 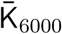, respectively (Supplementary Fig. 5). To simulate the polymer dynamics during chromosome condensation, we linearly interpolated the matrices 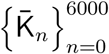 at 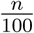 min as follows: 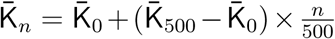 for 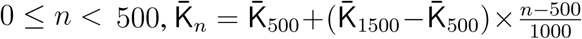 for 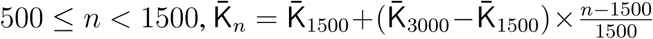 for 1500 *≤ n <* 3000, and 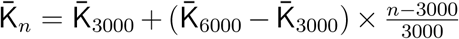 for 3000 *≤ n ≤* 6000. By using 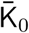, an initial polymer conformation was sampled. Then, the polymer dynamics between 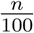 min and 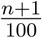 min was calculated by 1000 steps of numerical integration with 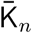 based on the integral algorithm (equations (1) and (2)).

In the visualization, we fixed the centre-of-mass of polymer conformations to the origin (Fig. 2f, Supplementary Videos 4, 5).

### Calculating shape length of polymer conformation

To quantify the characteristic shape of a polymer conformation 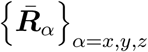 during chromosome condensation, we evaluated the characteristic shape lengths as an ellipsoidal conformation based on the gyration tensor 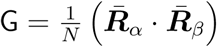. We calculated the three eigenvalues 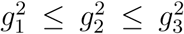 of the tensor G. Then, we adopted *g*_3_ and 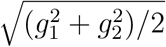 as the characteristic shape lengths of the major and minor axes, respectively (Fig. 2g).

## Code availability

Python codes of PHi-C are available at https://github.com/soyashinkai/PHi-C/Codes. Scripts to generate data and the figures shown in Fig. 1a can be found at https://github.com/soyashinkai/PHi-C/Demos.

## Data availability

Published publicly available Hi-C data were used in this study: Ohno et al. ^14^ (PRJNA427106), Rao et al. ^10^ (GSE63525), Bonev et al. ^16^ (GSE96107), and Gibcus et al. ^18^ (GSE102740). Input Hi-C matrix data of PHi-C were generated through the JUICER and JUICER TOOLS ^12^.

