## Supplementary Information for "PHi-C: deciphering Hi-C data into polymer dynamics"

#### Supplementary Note

Defining contacts in polymer modelling

Contact probability of fractal polymers

Theory of polymer network model

#### Supplementary Figures

Supplementary Figure 1

Supplementary Figure 2

Supplementary Figure 3

Supplementary Figure 4

Supplementary Figure 5

#### Supplementary Videos

Supplementary Video 1

Supplementary Video 2

Supplementary Video 3

Supplementary Video 4

Supplementary Video 5

#### Supplementary Note

**Defining contacts in polymer modelling.** The bead-spring model in polymer physics consists of $N$ beads (or monomers) connected by $N-1$ harmonic springs along the polymer backbone (Fig. 1b), where a polymer conformation is represented by $\{\boldsymbol{R}_i = (R_{i,x}, R_{i,y}, R_{i,z})\}_{i=0}^{N-1}$ and the characteristic length between adjacent monomers is expressed by $b$. The linearity of the harmonic springs enables us to analytically calculate the probability of the conformation $\boldsymbol{r}_{ij} = \boldsymbol{R}_i - \boldsymbol{R}_j$ between the $i$-th and $j$-th monomers as follows: $p(\boldsymbol{r}_{ij})\, d\boldsymbol{r}_{ij} = (2\pi\Sigma_{ij}^2)^{-\frac{3}{2}}\, e^{-\frac{x^2+y^2+z^2}{2\Sigma_{ij}^2}}\, dxdydz$, where $\Sigma_{ij}^2 = \left\langle \boldsymbol{r}_{ij}^2 \right\rangle /3$ is the variance for each freedom. Then, the probability density function for the distance $r_{ij} = |\boldsymbol{r}_{ij}|$ becomes $p(r_{ij}) = \sqrt{\frac{2}{\pi}}\,\Sigma_{ij}^{-3}\, r_{ij}^2\, e^{-\frac{r_{ij}^2}{2\Sigma_{ij}^2}}$. So far, a mathematical expression to define contacts within the contact distance $\sigma$ has been written by $\int_0^\sigma p(r_{ij})\, dr_{ij}$ [6,24]. However, the expression makes it difficult to improve further analytical calculations. Therefore, we introduced the contact Gaussian kernel with $\sigma$ (Fig. 1b). Then, the contact probability between the $i$-th and $j$-th monomers is represented by $C_{ij} \propto \int_0^\infty p(r_{ij}) e^{-\frac{r_{ij}^2}{2\sigma^2}}\, dr_{ij}$. Finally, under the normalization $C_{ii} = 1$, the contact probability can be expressed as

$$C_{ij} = \left(1 + \frac{\Sigma_{ij}^2}{\sigma^2}\right)^{-3/2}, \qquad \text{(S1)}$$

which means that the ratio of the conformational fluctuation $\Sigma_{ij}$ to the contact distance $\sigma$ is a physically important parameter to define the contacts. Note that equation (S1) was also recently derived with a similar consideration [25].

**Contact probability of fractal polymers.** The fractal polymer with the fractal dimension $d_{\mathrm{f}}$ is a generalized model of the bead-spring model with the scaling $\Sigma_{ij}^2 = \frac{b^2}{3}|i-j|^{2/d_{\mathrm{f}}}$, and the fractal dimension can characterize polymer condensation states [13,26] (Supplementary Fig. 2a). Substituting the scaling into equation (S1), we obtained a mathematical relation for the contact probability as a function of genomic distance $s$,

$$P(s) = \left(1 + \frac{1}{3(\sigma/b)^2}\left(\frac{s}{c}\right)^{2/d_\mathrm{f}}\right)^{-3/2}, \tag{S2}$$

where $c$ is the genomic size corresponding to a modelled monomer. Theoretical curves for the decay show two features (Supplementary Fig. 2b): (i) the ratio $\sigma/b$ results in a rounded shape at a small genomic distance, and (ii) the fractal dimension determines the scaling at a large genomic distance,

$$P(s) \sim s^{-3/d_\mathrm{f}}. \tag{S3}$$

These equations (S2) and (S3) allow us to extract physical information, $\sigma/b$ and $d_\mathrm{f}$, from experimental $P(s)$ data by fitting the parameters.

**Theory of polymer network model.** The polymer network model is a generalized model of the bead-spring model [27], and is characterized by the interaction matrix $\mathsf{K} = (k_{ij})$, which is an $N \times N$ matrix for an $N$-monomers polymer system. The element $k_{ij}$ generally represents the intensity of attractive or repulsive interaction between the $i$-th and $j$-th monomers. Therefore, the matrix satisfies $k_{ii} = 0$ and $\mathsf{K} = \mathsf{K}^\mathsf{T}$. Basically, the elastic force between two monomers results from an attractive interaction with a positive value. Furthermore, here, we take into account negative values, which represent repulsive interactions between monomers, as long as the system is stable as stated below.

The dynamics of the system $\{\boldsymbol{R}_0, \boldsymbol{R}_1, \cdots, \boldsymbol{R}_{N-1}\}$ is described by the Langevin equation

$$\gamma \frac{d\boldsymbol{R}_i(t)}{dt} = \sum_{j=0}^{N-1} k_{ij} \left[\boldsymbol{R}_j(t) - \boldsymbol{R}_i(t)\right] + \boldsymbol{g}_i(t), \tag{S4}$$

where $\gamma$ is the friction coefficient of the monomers. The thermal random force $\boldsymbol{g}_i(t)$ satisfies zero average, $\langle g_{i,\alpha}(t)\rangle = 0$, and the fluctuation-dissipation relation (FDR):

$$\langle g_{i,\alpha}(t) g_{j,\beta}(s)\rangle = 2\gamma k_{\mathrm{B}} T \delta_{\alpha\beta}\delta_{ij}\delta(t-s), \tag{S5}$$

where $k_{\mathrm{B}}$ is the Boltzmann constant, $T$ is the temperature of the environment, $\langle\cdot\rangle$ stands for the averaging at thermal equilibrium, and the suffixes $\alpha$ and $\beta$ represent $x$, $y$ and $z$.

According to graph theory [28], the matrix $\mathsf{K}$ can be converted into the Laplacian matrix $\mathsf{L}$ to characterize the properties of the network. The degree matrix is defined by

$$\mathsf{D} = \mathrm{diag}(D_0, D_1, \cdots, D_{N-1}), \tag{S6}$$

where

$$D_i = \sum_{j=0}^{N-1} k_{ij}, \tag{S7}$$

and the Laplacian matrix is derived as

$$\mathsf{L} = (L_{ij}) = \mathsf{D} - \mathsf{K}. \tag{S8}$$

Then, using the matrix $\mathsf{L}$, equation (S4) is rewritten as

$$\gamma \frac{d\boldsymbol{R}_i(t)}{dt} = -\sum_{j=0}^{N-1} L_{ij}\boldsymbol{R}_j(t) + \boldsymbol{g}_i(t). \tag{S9}$$

Finally, for the $\alpha$ coordinate vector of the monomers $\boldsymbol{R}_\alpha = (R_{0,\alpha}(t), R_{1,\alpha}(t), \cdots, R_{N-1,\alpha}(t))^{\mathsf{T}}$, the system is concisely described by

$$\gamma \frac{d\boldsymbol{R}_\alpha}{dt} = -\mathsf{L}\boldsymbol{R}_\alpha + \boldsymbol{g}_\alpha, \tag{S10}$$

where $\boldsymbol{g}_\alpha$ represents the vector $(g_{0,\alpha}(t), g_{1,\alpha}(t), \cdots, g_{N-1,\alpha}(t))^\mathsf{T}$.

As defined by equation (S8), the matrix $\mathsf{L}$ is real symmetric. Thus, $\mathsf{L}$ is diagonalizable. Furthermore, the $N$ eigenvalues are non-negative, so that the system is stable and the minimal eigenvalue is unique with a value of 0 due to the connectivity of the polymer network. Therefore, the $N$ eigenvalues satisfy

$$0 = \lambda_0 < \lambda_1 \leq \lambda_2 \leq \cdots \leq \lambda_{N-1}, \tag{S11}$$

and there is an orthogonal matrix $\mathsf{Q}$ such that

$$\mathsf{Q}^\mathsf{T}\mathsf{L}\mathsf{Q} = \mathrm{diag}(\lambda_0, \lambda_1, \cdots, \lambda_{N-1}) = \mathsf{\Lambda}. \tag{S12}$$

Equation (S8) implies that the orthonormal eigenvector $\boldsymbol{v}_0$ corresponding to $\lambda_0 = 0$ is proportional to $(1, 1, \cdots, 1)^\mathsf{T}$. That is, $\boldsymbol{v}_0 = (1/\sqrt{N}, 1/\sqrt{N}, \cdots, 1/\sqrt{N})^\mathsf{T}$. Using the other orthonormal eigenvectors $\boldsymbol{v}_p$ corresponding to $\lambda_p$, the orthogonal matrix is expressed as

$$\mathsf{Q} = \left((1/\sqrt{N}, 1/\sqrt{N}, \cdots, 1/\sqrt{N})^\mathsf{T}\, \boldsymbol{v}_1\, \boldsymbol{v}_2 \cdots \boldsymbol{v}_{N-1}\right). \tag{S13}$$

The matrix $\mathsf{L}$ is singular matrix due to $\lambda_0 = 0$. However, introducing a diagonal matrix

$$\tilde{\mathsf{\Lambda}}^{-1} = \mathrm{diag}(0, \lambda_1^{-1}, \lambda_2^{-1}, \cdots, \lambda_{N-1}^{-1}), \tag{S14}$$

we can obtain a matrix

$$\mathsf{M} = (M_{ij}) = \mathsf{Q}\tilde{\mathsf{\Lambda}}^{-1}\mathsf{Q}^\mathsf{T} = \left(\sum_{p=1}^{N-1} Q_{ip}Q_{jp}\lambda_p^{-1}\right), \tag{S15}$$

which satisfies the following relation:

$$\mathsf{LM} = \mathsf{I} - \begin{pmatrix} 1/N & 1/N & \dots & 1/N \\ 1/N & 1/N & \dots & 1/N \\ \vdots & \vdots & \ddots & \vdots \\ 1/N & 1/N & \dots & 1/N \end{pmatrix}. \tag{S16}$$

The physical implication of the matrix $\mathsf{M}$ is revealed below in equation (S21).

Multiplying the matrix $\mathsf{Q}^\mathsf{T}$ to equation (S10) from the left, the dynamics for the transformed vector $\boldsymbol{X}_\alpha = \mathsf{Q}^\mathsf{T}\boldsymbol{R}_\alpha = (X_{0,\alpha}(t), X_{1,\alpha}(t), \cdots, X_{N-1,\alpha}(t))$ obeys

$$\gamma\frac{d\boldsymbol{X}_\alpha}{dt} = -\Lambda\boldsymbol{X}_\alpha + \mathsf{Q}^\mathsf{T}\boldsymbol{g}_\alpha \tag{S17}$$

with the FDR (equation (S5)). This equation represents the orthogonalized linear Langevin equation for the polymer network model, and the stochastic differential equation reveals the Ornstein–Uhlenbeck process [29]. Thus, the covariance for the vector $\boldsymbol{X}_\alpha$ satisfies the following relation:

$$\langle X_{p,\alpha}X_{q,\beta}\rangle = \frac{k_\mathrm{B}T}{\lambda_p}\delta_{pq}\delta_{\alpha\beta} \quad (p, q = 1, , 2, \cdots, N-1). \tag{S18}$$

According to the form of the orthogonal matrix $\mathsf{Q}$ in equation (S13), the centre-of-mass of a polymer conformation is expressed by

$$\frac{X_{0,\alpha}}{\sqrt{N}} = \frac{1}{\sqrt{N}}\sum_{i=0}^{N-1}\left(\mathsf{Q}^\mathsf{T}\right)_{0i}R_{i,\alpha} = \frac{1}{N}\sum_{i=0}^{N-1}R_{i,\alpha}. \tag{S19}$$

Then, the transformation for the original coordinates, $\boldsymbol{R}_\alpha = \mathsf{Q}\boldsymbol{X}_\alpha$, corresponds to

$$R_{i,\alpha} - \frac{X_{0,\alpha}}{\sqrt{N}} = \sum_{p=1}^{N-1}Q_{ip}X_{p,\alpha}. \tag{S20}$$

Therefore, the covariance between the $i$-th and $j$-th monomers with respect to the centre-of-mass is calculated as follows:

$$\begin{aligned}
\left\langle \left[R_{i,\alpha} - \frac{X_{0,\alpha}}{\sqrt{N}}\right]\left[R_{j,\beta} - \frac{X_{0,\beta}}{\sqrt{N}}\right]\right\rangle &= \left\langle \left[\sum_{p=1}^{N-1} Q_{ip} X_{p,\alpha}\right]\left[\sum_{q=1}^{N-1} Q_{jq} X_{q,\beta}\right]\right\rangle \\
&= \sum_{p=1}^{N-1}\sum_{q=1}^{N-1} Q_{ip} Q_{jq} \frac{k_{\mathrm{B}}T}{\lambda_p}\delta_{pq}\delta_{\alpha\beta} \\
&= k_{\mathrm{B}}T\delta_{\alpha\beta}\sum_{p=1}^{N-1} Q_{ip}Q_{jp}\lambda_p^{-1} \\
&= k_{\mathrm{B}}T\delta_{\alpha\beta}M_{ij}. \qquad \text{(S21)}
\end{aligned}$$

Here, we used equations (S15) and (S20). Thus, we can understand that the matrix $\mathsf{M}$ represents the covariance matrix between positions of the $i$-th and $j$-th monomers relative to the centre-of-mass.

To calculate the contact matrix $\mathsf{C} = (C_{ij})$ based on equation (S1), we should know the variance matrix $\Sigma^2 = \left(\Sigma_{ij}^2\right)$ between the $i$-th and $j$-th monomers for each freedom. The calculations are as follows:

$$\begin{aligned}
\Sigma_{ij}^2 &= \left\langle [R_{i,\alpha} - R_{j,\alpha}]^2 \right\rangle \\
&= \left\langle \left[\left(R_{i,\alpha} - \frac{X_{0,\alpha}}{\sqrt{N}}\right) - \left(R_{j,\alpha} - \frac{X_{0,\alpha}}{\sqrt{N}}\right)\right]^2 \right\rangle \\
&= \left\langle \left[R_{i,\alpha} - \frac{X_{0,\alpha}}{\sqrt{N}}\right]^2 \right\rangle + \left\langle \left[R_{j,\alpha} - \frac{X_{0,\alpha}}{\sqrt{N}}\right]^2 \right\rangle - 2\left\langle \left[R_{i,\alpha} - \frac{X_{0,\alpha}}{\sqrt{N}}\right]\left[R_{j,\alpha} - \frac{X_{0,\alpha}}{\sqrt{N}}\right]\right\rangle \\
&= k_{\mathrm{B}}T\left[M_{ii} + M_{jj} - 2M_{ij}\right]. \qquad \text{(S22)}
\end{aligned}$$

Note that the matrix transformation between $\mathsf{M}$ and $\Sigma^2$ is invertible because the matrix $\mathsf{M}$ satisfies an additional condition $\sum_j M_{ij} = 0$.

The mean-squared displacement (MSD) of the $i$-th monomer is analytically calculated as follows:

$$\begin{aligned}
\mathrm{MSD}(t;i) &= \left\langle \sum_{\alpha=x,y,z} \left[R_{i,\alpha}(t) - R_{i,\alpha}(0)\right]^2 \right\rangle \\
&= \left\langle \sum_{\alpha=x,y,z} \left[ \left( \frac{X_{0,\alpha}(t)}{\sqrt{N}} - \frac{X_{0,\alpha}(0)}{\sqrt{N}} \right) + \sum_{p=1}^{N-1} Q_{ip} \left( X_{p,\alpha}(t) - X_{p,\alpha}(0) \right) \right]^2 \right\rangle \\
&= 3 \left\langle \left[ \left( \frac{X_0(t)}{\sqrt{N}} - \frac{X_0(0)}{\sqrt{N}} \right) + \sum_{p=1}^{N-1} Q_{ip} \left( X_p(t) - X_p(0) \right) \right]^2 \right\rangle \\
&= 3 \left\langle \left( \frac{X_0(t)}{\sqrt{N}} - \frac{X_0(0)}{\sqrt{N}} \right)^2 \right\rangle + 6 \sum_{p=1}^{N-1} Q_{ip}^2 \left( \left\langle X_p^2 \right\rangle - \left\langle X_p(t) X_p(0) \right\rangle \right), \qquad \text{(S23)}
\end{aligned}$$

where we used equation (S20) and the spatial isotropy. The first term of the right-hand side corresponds to the MSD of the centre-of-mass and is written as [13]

$$\frac{6k_{\mathrm{B}}T}{N\gamma}\, t. \qquad \text{(S24)}$$

The Langevin dynamics for the transformed vector $\boldsymbol{X}_p$ (equation (S17)) and the variance relation (equation (S18)) derive that the autocorrelation function decays exponentially,

$$\begin{aligned}
\left\langle X_p(t) X_p(0) \right\rangle &= \left\langle X_p^2 \right\rangle e^{-\lambda_p/\gamma \cdot t} \\
&= k_{\mathrm{B}} T \lambda_p^{-1} e^{-\lambda_p/\gamma \cdot t}. \qquad \text{(S25)}
\end{aligned}$$

Finally, the MSD obeys

$$\mathrm{MSD}(t;i) = \frac{6k_{\mathrm{B}}T}{N\gamma}\, t + 6k_{\mathrm{B}}T \sum_{p=1}^{N-1} Q_{ip}^2 \lambda_p^{-1} \left( 1 - e^{-\lambda_p/\gamma \cdot t} \right). \qquad \text{(S26)}$$

#### Supplementary Figures

**Supplementary Figure 1: Algorithms of PHi-C. a**, Normalization procedure for PHi-C input data. Input data are a contact matrix $C_{ij(\mathrm{JUICER})}$ generated through the JUICER and JUICER TOOLS [12]. First, the contact probability $P_{\mathrm{JUICER}}(|i-j|)$ is calculated by averaging $C_{ij(\mathrm{JUICER})}$. If there is no value of $P_{\mathrm{JUICER}}(|i-j|)$ due to a lack of read counts, one must interpolate an appropriate value so that the shape of the contact probability is consistent. Next, $C_{ij(\mathrm{JUICER})}$ with a value of zero is interpolated by $P_{\mathrm{JUICER}}(|i-j|)$, and $C_{ij(\mathrm{interpolated})}$ is obtained. In this interpolation, the contact probability $P_{\mathrm{JUICER}}(|i-j|)$ is unaltered. Finally, dividing $C_{ij(\mathrm{interpolated})}$ by $\langle C_{ii}\rangle = P_{\mathrm{JUICER}}(0)$, a normalized Hi-C contact matrix $\mathsf{C}_{\mathsf{Hi-C}}$ is obtained. In the double-logarithmic plot, the shape of the normalized contact probability is unchanged. **b**, Flowchart of the optimization algorithm. **c**, Assessing the optimization algorithm of PHi-C. First, a normalized interaction matrix $\bar{\mathsf{K}}$ was randomly generated. Second, $\bar{\mathsf{K}}$ was converted into a contact matrix $\mathsf{C}_{\mathsf{generated}}$ (upper) as input data for optimization. Next, through running optimization, an optimized contact matrix $\mathsf{C}_{\mathsf{optimized}}$ (lower) was obtained. Finally, Pearson's correlation coefficient $r$ between $\log_{10}\mathsf{C}_{\mathsf{optimized}}$ and $\log_{10}\mathsf{C}_{\mathsf{generated}}$ was calculated for validation. The histogram for 100 randomly generated samples shows that the optimization of PHi-C provides more than 0.998 correlation. The best (right; $r = 0.999975$) and worst (left; $r = 0.998536$) samples are shown. Here, the matrix size is $100 \times 100$.

**Supplementary Figure 2: Fractal organization in contact probabilities.** **a**, Fractal polymer model. The fractal dimension $d_\mathrm{f}$ characterize polymer condensation states [13,26]. **b**, Theoretical curves of the contact probability $P(s)$ in equation (3). $P(s)$ is normalized by the value at $s/c = |i-j| = 1$, where $i$ and $j$ are monomer indices of the fractal polymer and $c$ represents the genomic size corresponding to every monomer. (Left) $P(s)$ for fixed $d_\mathrm{f} = 2.0$, and $\sigma/b = 0.2,\ 0.5,\ 1.0,\ 2.0$ and $5.0$. (Right) $P(s)$ for fixed $\sigma/b = 1.0$, and $d_\mathrm{f} = 1.5,\ 2.0,\ 3.0$ and $4.0$. Theoretical scaling relations, as in equation (4), are shown. **c**, Averaged contact probability $P(s)$ for the yeast Hi-C data [14] with 160-bp resolution. The fitted values, $\sigma/b$ and $d_\mathrm{f}$, by using equations (3) and (4) are displayed. The arrow shows a characteristic genomic distance: where two fitted curves cross, this indicates the hierarchically fractal organization. **d**, Averaged contact probability $P(s)$ for the GM12878 Hi-C data in situ primary [10]. $P(s)$ was calculated with 1-kb resolution for the 2–20-kb region, 10-kb resolution for the 20–200-kb region, and 100-kb resolution for the 200-kb–50-Mb region. The values fitted by using equations (3) and (4) are displayed. The arrows show characteristic genomic distances where two fitted curves cross, indicating the hierarchically fractal organization.

**Supplementary Figure 3: Dynamic feature of *Nanog* and *Oct4* loci in mESCs. a**, TAMSDs of the two loci within 300 s. Thick curves represent the ensemble-averaged TAMSDs of *Nanog* and *Oct4* for 29 and 32 trajectories, respectively. The coloured area represents the standard deviation. Beyond 175 s, the *Nanog* movement revealed a statistically significant enhancement compared to the *Oct4* movement. **b**, All TAMSD data within 490 s in double-logarithmic scale. (Upper) Thick curves represent the ensemble-averaged TAMSDs. Within 300 s, 29 and 32 TAMSDs of *Nanog* and *Oct4* are sub-diffusive (slope $< 1$), respectively. A theoretical consideration regarding polymer physics has indicated that the intra-chromosome movement results in sub-diffusion [13]. We adopted these TAMSDs for further analysis. (Lower) TAMSDs for the other trajectories indicate normal diffusion (slope $= 1$), in which other factors should drive the normal-diffusive chromosome movement. We eliminated these TAMSDs. **c**, Adopted TAMSDs within 300 s. (Upper) Thick curves represent the ensemble-averaged TAMSDs. (Lower) Time series of $p$-value of Welch's $t$-test. Beyond 175 s, the $p$-values are less than 0.07.

**Supplementary Figure 4: Optimized contact probability data for chromosome 7 of DT40 cells at G2 (0 min), 5, 15, 30, and 60 min. a**, Correlation plots between $\log_{10} \mathsf{C}_{\mathsf{Hi-C}}$ and $\log_{10} \mathsf{C}_{\mathsf{Optimized}}$. **b**, Contact probabilities as a function of genomic distance $s$.

**Supplementary Figure 5: Optimized interaction parameters in the polymer network model for chromosome 7 of DT40 cells at G2 (0 min), 5, 15, 30, and 60 min.** (Upper) Optimized interaction matrices $\frac{\sigma^2}{3k_{\mathrm{B}}T}\left(k_{ij}\right) = \left(\bar{k}_{ij}\right)$. (Lower) Optimized interaction parameters along the polymer backbone $\frac{\sigma^2}{3k_{\mathrm{B}}T}\, k_{i,i+1} = \bar{k}_{i,i+1}$.

#### Supplementary Videos

**Supplementary Video 1** (Left) A 4D simulation, where $10^6$ steps of numerical integration were carried out, of the polymer network model with intra- and inter domain interactions in Fig. 1e. The polymer is coloured according to the bottom figure in Fig. 1e. (Right) The corresponding 3D distance map.

**Supplementary Video 2** (Left) A 4D simulation, where $10^6$ steps of numerical integration were carried out, of the polymer network model with loop interactions in Fig. 1f. The polymer is coloured according to the bottom figure in Fig. 1f. (Right) The corresponding 3D distance map.

**Supplementary Video 3** (Left) A 4D simulation, where $10^6$ steps of numerical integration were carried out, of the polymer network model with heterogeneous connectivity along the polymer backbone in Fig. 1g. The polymer is coloured according to the bottom figure in Fig. 1g. (Right) The corresponding 3D distance map.

**Supplementary Video 4** (Left) A 4D simulation, where $6 \times 10^6$ steps of numerical integration were carried out, during mitotic chromosome formation of chromosome 7 in chicken DT-40 cells. (Right) Optimized contact matrix $\{\mathsf{C}_n\}_{n=0}^{6000}$ at $\frac{n}{100}$ min. Through the optimization of PHi-C, we obtained the optimized matrices $\bar{\mathsf{K}}_0$, $\bar{\mathsf{K}}_{500}$, $\bar{\mathsf{K}}_{1500}$, $\bar{\mathsf{K}}_{3000}$ and $\bar{\mathsf{K}}_{6000}$. Then we generated $\left\{\bar{\mathsf{K}}_n\right\}_{n=0}^{6000}$ with linear interpolation. Finally, we transformed $\bar{\mathsf{K}}_n$ into $\mathsf{C}_n$.

**Supplementary Video 5** Spinning 3D structures at G2 (0 min), 5, 15, 30 and 60 min in the 4D simulation (Supplementary Video 4).

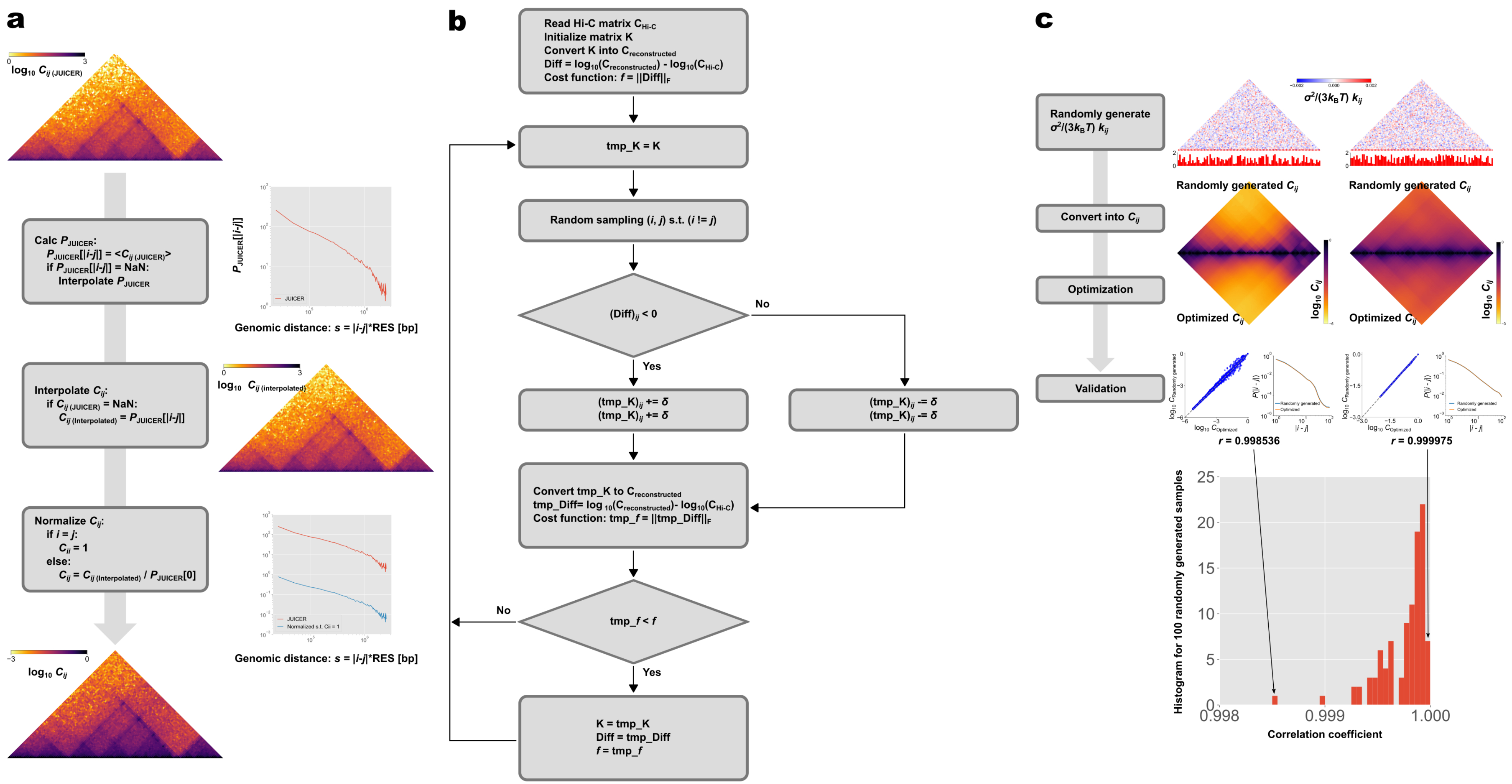


### Supplementary Figure 1

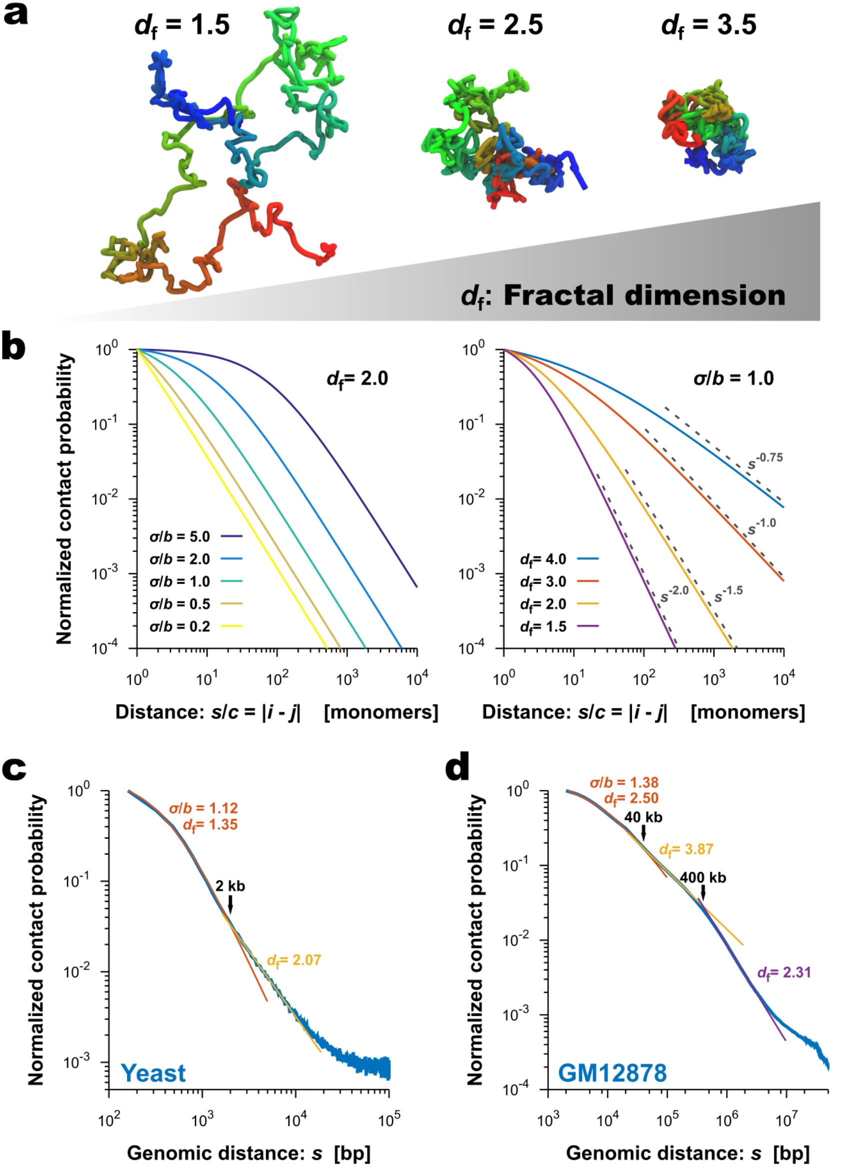


### Supplementary Figure 2

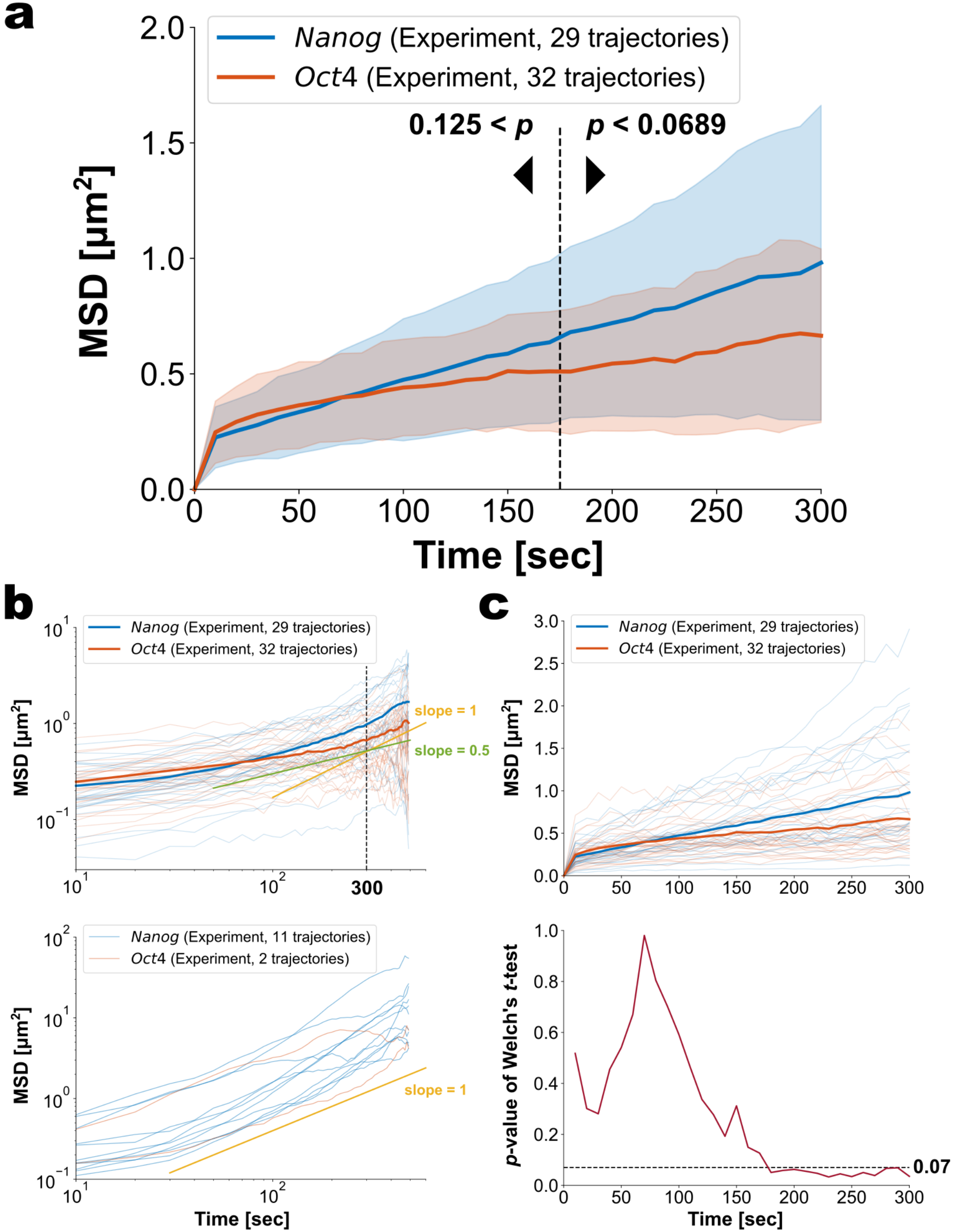


### Supplementary Figure 3

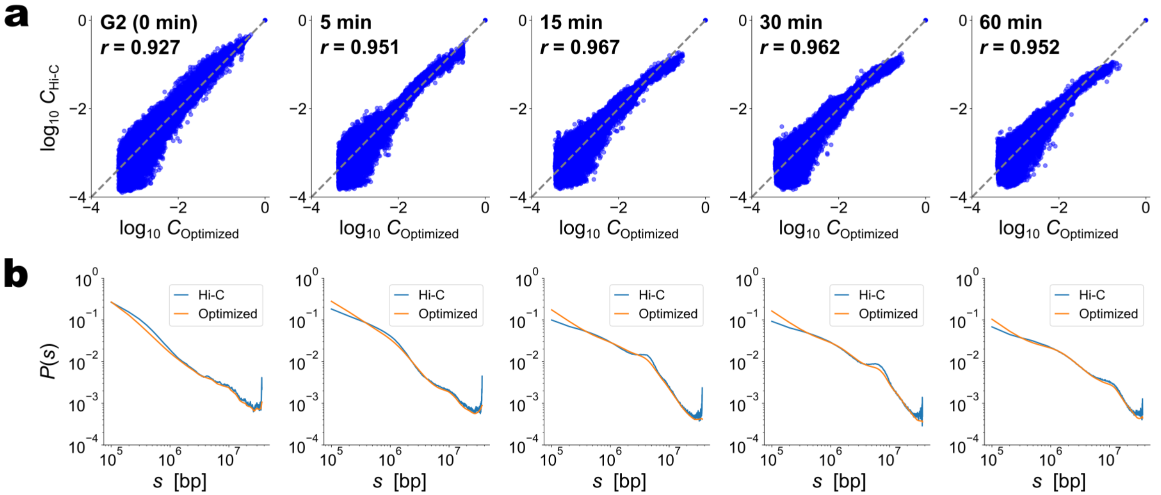


### Supplementary Figure 4

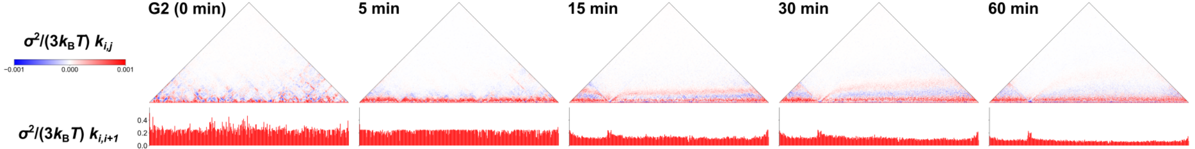


### Supplementary Figure 5
